# Environment determines evolutionary trajectory in a constrained phenotypic space

**DOI:** 10.1101/096909

**Authors:** David T. Fraebel, Harry Mickalide, Diane Schnitkey, Jason Merritt, Thomas E Kuhlman, Seppe Kuehn

## Abstract

Constraints on phenotypic variation limit the capacity of organisms to adapt to the multiple selection pressures encountered in natural environments. To better understand evolutionary dynamics in this context, we select *Escherichia coli* for faster migration through a porous environment, a process which depends on both motility and growth. We find that a trade-off between swimming speed and growth rate constrains the evolution of faster migration. Evolving faster migration in rich medium results in slow growth and fast swimming, while evolution in minimal medium results in fast growth and slow swimming. In each condition parallel genomic evolution drives adaptation through different mutations. We show that the trade-off is mediated by antagonistic pleiotropy through mutations that affect negative regulation. A model of the evolutionary process shows that the genetic capacity of an organism to vary traits can qualitatively alter the evolutionary trajectory when selection pressures are complex.

## Introduction

In Nature organisms adapt to complex environments where many biotic and abiotic factors affect survival. For microbes these factors include demands on metabolism (*Savageau, 1983*), motility (*Celani and Vergassola, 2010*) and antibiotic resistance (*Vetsigian et al., 2011*). In this context, evolution involves the simultaneous adaptation of many phenotypic traits. Organisms under complex selection pressures often cannot vary traits independently and instead exhibit trade-offs (*Shoval et al., 2012*).

Trade-offs constrain adaptive responses to selection. For example, phage exhibit a trade-off between fecundity and virulence which depends on the relative duration of periods of horizontal and vertical transmission (*Messenger et al., 1999*). Bacterial populations selected for efficient conversion of nutrients to biomass exhibit a trade-off between yield and growth rate (*Bachmann et al., 2013*).

Predicting evolution in complex environments requires quantifying both trade-offs and selection pressures (*Lande, 1979*). In wild populations of birds (*Grant and Grant, 1995*) and fish (*Ghalambor et al., 2003*), phenotypic constraints and selection pressures have been inferred from measurements of phenotypic variation. However, in wild populations of higher organisms it is challenging to observe evolution, determine selection pressures and elucidate mechanisms constraining phenotypes. To better understand the interplay between trade-offs, selection and evolution it is necessary to study genetically tractable, rapidly evolving microbial populations in the laboratory.

However, laboratory based experimental evolution of microbes typically selects for a single phenotype such as growth rate (*Lang et al., 2013*). There is evidence that metabolic trade-offs arise in these experiments from the decay of traits that are not subject to selection (*Cooper and Lenski, 2000*) rather than a compromise between multiple selection pressures. Other experiments explore how phenotypes restricted by trade-offs evolve under alternating selection for individual traits (*Yi and Dean, 2016; Messenger et al., 1999*). Less is known about evolutionary dynamics in the naturally relevant regime where selection pressures are multifaceted.

To address this, we selected *Escherichia coli* for faster migration through a porous environment. We showed that the evolution of faster migration is constrained by a trade-off between swimming speed and growth rate. Evolution of faster migration in rich medium is driven by faster swimming despite slower growth, while faster migration in minimal medium is achieved through faster growth despite slower swimming. Sequencing and genetic engineering showed that this trade-off is due to antagonistic pleiotropy through mutations that affect negative regulation. Finally, a model of multi-trait selection supports the hypothesis that the direction of evolution along the Pareto frontier that constrains accessible phenotypes is determined by the genetic variance of each trait. Our results show that when selection acts simultaneously on two traits governed by a trade-off, the environment determines the evolutionary trajectory.

## Results

### Experimental evolution of migration rate

E. *coli* inoculated at the center of a low viscosity agar plate consume nutrients locally, creating a spatial nutrient gradient which drives chemotaxis through the porous agar matrix (*Righetti et al., 1981; Maaloum et al., 1998*) and subsequent nutrient consumption (*Adler, 1966; Wolfe and Berg, 1989; Croze et al., 2011*). The result is a three-dimensional bacterial colony that expands radially across the plate as individuals swim and divide in the porous environment. We refer to the outermost edge of an expanding colony as the migrating front. We tracked these migrating fronts using webcams and light-emitting diode (LED) illumination (Methods). The front migrates at a constant speed *_s_* after an initial growth phase (*Adler, 1966; Wolfe and Berg, 1989*).

We performed experimental evolution by repeating rounds of allowing a colony to expand for a fixed interval, selecting a small population of cells from the migrating front and using them to inoculate a fresh low viscosity agar plate (Figure 1(a)). We performed selection experiments in this way for two distinct nutrient conditions. First, we used rich medium (lysogeny broth (LB), 0.3 % w/v agar, 30°C) where all amino acids are available. In this medium the population forms concentric rings (Figure 1(b)) that consume amino acids sequentially. The outermost ring consumes L-serine and most of the oxygen (*Adler, 1966*). Second, we used minimal medium (M63, 0.18mM galactose, 0.3 % w/v agar, 30°C) where populations migrate towards and metabolize galactose.

**Figure 1.**
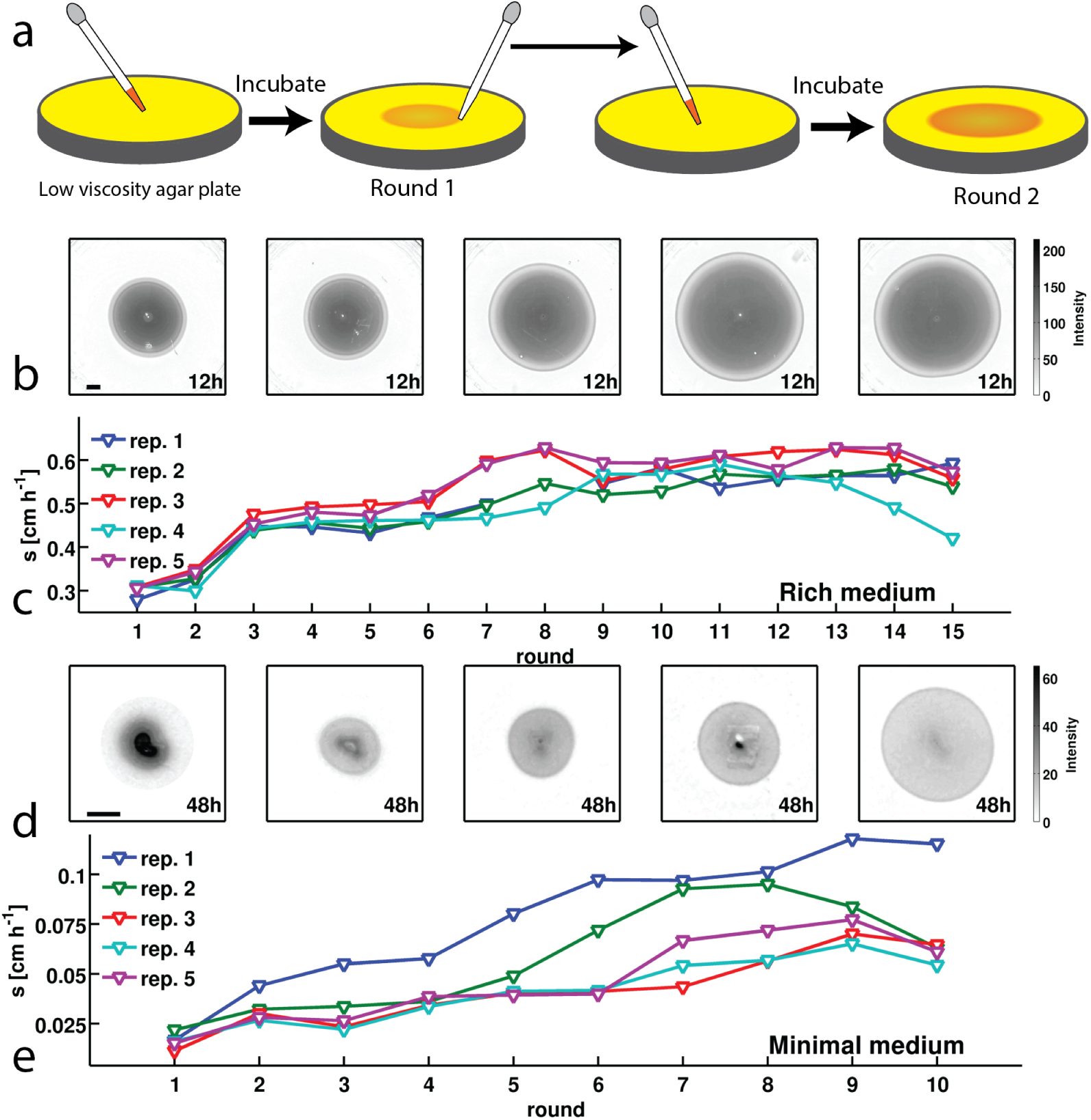
*E. coli* evolves faster migration through a porous environment in rich and minimal media: (a)A schematic of the selection procedure. *E. coli* are inoculated into the center ofa low viscosity (0.3 % w/v) agar plate where they form an expanding colony driven by metabolism and motility. After a fixed period of incubation a sample is taken from the outer edge of the expanded colony and used to inoculate a fresh plate. (b) Shows expanded colonies in rich medium (LB) plates after 12 hours of incubation over five successive rounds of selection. The color bar to the right applies to all panels in (b), with darker gray indicating higher cell density. Image intensity is assumed to be monotonic but not linear with cell density in the plate. Scale bar in the left panel is 1 cm and applies to all panels in (b). (c) Shows the rate of migration as a function of round of selection over 15 rounds for five replicate selection experiments in rich medium. (d) Shows colonies (gray regions) in minimal medium (M63, 0.18mM galactose) after 48 hours of incubation. The color bar to the right applies to all panels in (b). The scale bar in the left panel is 1 cm. (e) Shows the rate of migration as a function of round of selection over 10 rounds for five replicate selection experiments in minimal medium. Errors in migration rates were smaller than the size of markers. See Methods for details of image processing in both experiments. Figure 1- figure supplement 1: Selection with non-chemotactic (ΔcheA-Z) mutant, Figure 1 - figure supplement 2: Change in migration rate during long-term liquid culture, Figure 1 - figure supplement 3: Adaptation in rich medium depends on sampling location, Figure 1 - figure supplement 4: Comparison of founding and evolved strains to RP437: Single-cell swimming in rich medium, Figure 1 - figure supplement 5: Persistence of rich medium fast migrating phenotype in rich medium.

In rich medium colonies of wild-type bacteria (MG1655-motile, founding strain) expand with a front migration speed *s* ≈ 0.3 cm h^−1^ and cells were sampled from the front after 12 hours (Figure 1(b)). A portion of this sample was used to immediately inoculate a fresh plate while the remainder was preserved cryogenically. The process was repeated every 12 hours for 15 rounds. We observed a nearly two-fold increase in *s* over the course of the first 5 rounds of selection. The increase in *s* was reproducible across 5 independent selection experiments (Figure 1(c)).

To check whether chemotaxis was necessary for increasing *s*, we performed selection experiments using a motile but non-chemotactic mutant (Δ*cheA-Z*, Methods). Motility in this strain was confirmed by single-cell imaging in liquid media. As observed previously (*Wolfe and Berg, 1989*), the non-chemotactic strain formed dense colonies in low viscosity agar that remained localized near the site of inoculation and expanded ~1 cm in a 24 hour period: a rate 10-fold slower than the wild-type. To allow sufficient time for colony expansion, we performed selection experiments using this strain with 24 hour incubation times and observed an increase in *s* from approximately 0.03cm h^−1^ to 0.04cmh^−1^ (Figure 1 - figure supplement 1). We did not observe fast migrating spontaneous mutants which have been reported previously in multiple species (*Wolfe and Berg, 1989; Mohari et al., 2015*), likely because our plates were incubated for a shorter period of time.

To determine the number of generations transpiring in our selection experiments, we measured the number of cells in the inoculum and the number of cells in the colony after 12 hours of growth and expansion (Methods). We estimated that 10 to 12 generations occured in each round of selection. We then tested whether prolonged growth in well mixed liquid medium for a similar number of generations could lead to faster migration by growing the founding strain for 200 generations in continuous liquid culture and periodically inoculating a low viscosity agar plate (Figure 1 - figure supplement 2). We observed only a 3.5 % increase in the rate of migration, demonstrating that selection performed on spatially structured populations results in more rapid adaptation for fast migration than growth in well mixed conditions.

We then performed selection experiments in a minimal medium where growth and migration are substantially slower than in rich medium (Figure 1(d)). In this condition we allowed 48 hours for each round of expansion and took precautions to limit evaporative loss in the plates over this longer timescale (Methods). We observed an approximately 3-fold increase in *s* over the course of 10 rounds of selection, and this increase was reproduced across 5 replicate experiments. In the first round the population formed small ~1.5cm diameter colonies without a well defined front. Populations formed well defined fronts in subsequent rounds of selection (Figure 1(d)), reflecting a transition from growth and diffusion dominated transport to chemotaxis dominated migration (*Crozeetal., 2011*).

When we performed selection in minimal medium using the non-chemotactic mutant (∆*cheA-Z*), we found little or no migration and only a very small increase in the migration rate over 10 rounds of selection (Figure 1 - figure supplement 1). We concluded that chemotaxis is also necessary for increasing *s* in this medium.

Using the same technique described for rich medium, we estimated the number of generations per round of selection in minimal medium to be <10. We tested whether approximately 100 generations of growth in liquid was sufficient to evolve faster migration in minimal medium. Here we found that prolonged growth in well mixed conditions resulted in ~2-fold faster front migration. Despite the increase in migration rate, selection in well mixed conditions resulted in slower migration than selection in low viscosity agar plates for a similar number of generations (Figure 1 - figure supplement 2).

### Increasing swimming speed and growth rate increase migration rate

To characterize the adaptation we observed in Figure 1(c,e), we studied a reaction-diffusion model of migrating bacterial fronts of the type pioneered by Keller and Segel (*Keller and Segel, 1971*) and reviewed in Tindall *et al.* (*Tindall et al., 2008*). We model the bacterial density *ρ*(**r**,*t*) and a single chemo-attractant that also permits growth *c*(**r**,*t*). Our model includes only a single nutrient since the growth and chemotaxis of the outermost ring in rich media is driven by L-serine (*Adler, 1966*) and our minimal media conditions contain only a single carbon source/attractant. The dynamics of *ρ*(**r**,*t*) and *c*(**r**,*t*) are governed by

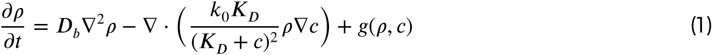

and

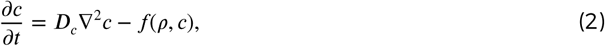

where the spatial and temporal dependence of *ρ* and *c* have been suppressed for clarity. The three terms on the right hand side of equation (1) describe diffusion, chemotaxis and growth respectively. ***D***_*b*_ is the bacterial diffusion constant, which describes the rate of diffusion of bacteria due to random, undirected motility. ***k***_0_ is the chemotactic coefficient capturing the strength of chemotaxis in response to gradients in attractant. ***K***_*D*_ is the equilibrium binding constant between the attractant and its associated receptor (*Brown and Berg, 1974*). Growth is modeled using the Monod equation 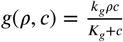, where *k_g_* is the maximum growth rate and ***K***_*g*_ is the half-maximum growth rate. *f*(*ρ,c*) describes the nutrient consumption and has an identical form to *g* since we assume the yield (***Y*** cells mL^−1^mM^−1^) is a constant. ***D***_*c*_ is the diffusion constant of small molecules in water. The physiological parameters describing growth and and attractant-receptor binding (*k_g_*,***K***_*g*_, ***Y*** and ***K***_*D*_) were either measured here or have been reported in the literature and can be applied directly in our simulation of migration in both nutrient conditions. Table 11 describes each parameter used in this study.

**Table 11.** Reaction-diffusion model parameters: Columns indicate parameter, explanation of parameter, units, value used in simulation of founder strain in rich medium, and the value used in simulation of founder strain in minimal medium. Parameters marked with an ^m^ were measured in this study. ***D***_*b*_, *k*_0_ and ***D***_*c*_ in rich medium were estimated as described in Supplementary file 1 using the methods of Croze *et al.* (2011). ***D***_*c*_ is assumed to be the same in minimal medium as rich medium. Identical *k*_0_ and ***D***_*b*_ were used in the minimal medium since Ford and Lauffenberger (1992) find nearly identical values for galactose as Ahmed and Stocker (2008) do for serine. ***K***_*D*_ for both nutrient conditions was taken from Adler, Hazelbauer and Dahl, (1973). For minimal medium ***K***_*g*_ and ***Y*** were taken from Lendenmann, Snozzi, and Egli (1999). The values cited for *s* were measured from numerical simulation of the reaction-diffusion model as outlined in Methods.

| parameter | explanation | units | founder value RM | founder value MM |
| --- | --- | --- | --- | --- |
| Single-cell swimming (this study) |  |  |  |  |
| $\tau_r$ | run duration | s | 0.67<sup>m</sup> | 0.47<sup>m</sup> |
| $\tau_t$ | tumble duration | s | – | – |
| $\vert v_r\vert $ | run speed | $\mu\text{m s}^{-1}$ | 18.7<sup>m</sup> | 22.2<sup>m</sup> |
| Reaction-diffusion model |  |  |  |  |
| $\rho(\mathbf{r}, t)$ | cell density | $\text{m}^{-3}$ | – | – |
| $c(\mathbf{r}, t)$ | nutrient density | mM | – | – |
| $c_0$ | nutrient concentration in medium | mM | 1<sup>m</sup> | 0.18<sup>m</sup> |
| $D_b$ | bacterial diffusion constant | $\text{cm}^2 \text{h}^{-1}$ | 0.0576 | 0.0576 |
| $D_c$ | nutrient diffusion constant | $\text{cm}^2 \text{h}^{-1}$ | 0.036 | 0.036 |
| $k_0$ | chemotactic coefficient in liquid | $\text{cm}^2 \text{h}^{-1}$ | 6.12 | 6.12 |
| $K_D$ | receptor-nutrient binding constant | mM | 2 | 0.1 |
| $k_g$ | maximum growth rate | $\text{h}^{-1}$ | 1.23<sup>m</sup> | 0.125<sup>m</sup> |
| $K_g$ | $c$ concentration for half-maximum growth rate | mM | 0.1 | $3 \times 10^{-4}$ |
| $Y$ | yield biomass per unit nutrients | cells<br>$\text{mL}^{-1} \text{mM}^{-1}$ | $5 \times 10^{7m}$ | $3 \times 10^8$ |
| $C$ | agar concentration | % (w/v) | 0.3<sup>m</sup> | 0.3<sup>m</sup> |
| $s$ | front migration rate | $\text{cm h}^{-1}$ | 0.61 | 0.09 |

The bacterial diffusion constant and the chemotactic coefficient depend on motility and the physical structure of the agar matrix. Motility in *E. coli* consists of runs, segments of nearly straight swimming ~0.5 to 1 second long at ~20μm^−1^, and tumbles that rapidly reorient the cell over a period of ~0.1 seconds (*Berg and Brown, 1972*). Rivero et al. showed how the reaction-diffusion parameters ***D***_*b*_ and *k*_0_ depend on run speed and duration (*Rivero et al., 1989*). Croze et al. (*Croze et al., 2011*) modified these results to account for the presence of the agar matrix. The approach treats interactions between cells and agar as scattering events where the cell is forced to tumble. See the supplementary file 1 for a detailed discussion of these models. We estimated ***D***_*b*_ and *k*_0_ using the method developed by Croze et al. for our conditions. With these parameters we simulated the model in equations (1) and (2) with parameters appropriate for rich media (chemotaxis towards L-serine) and minimal media (chemotaxis towards galactose) and found that it recapitulated the qualitative features of the migration observed experimentally (Figure 2 - figure supplement 1 and 2). The rate of front migration in our simulations was 0.61 cmh^−1^ for rich media and 0.09cmh^−1^ for minimal media. We note that this comparison involves no free parameters.

**Figure 2.**
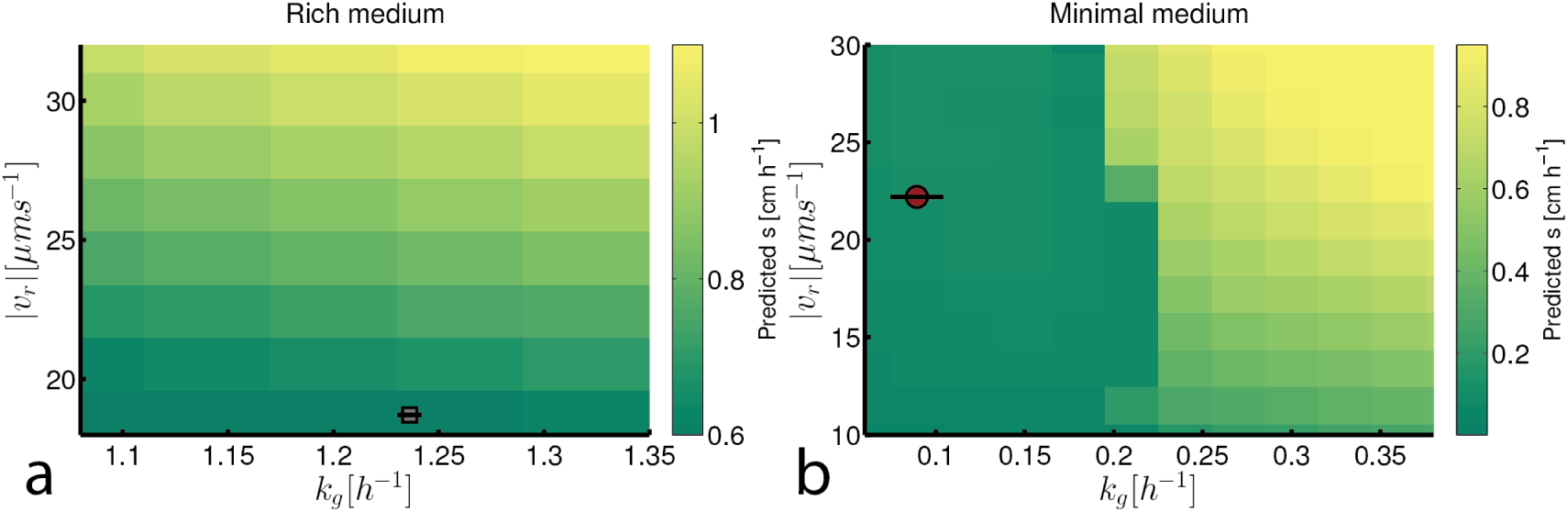
Migration rate increases with run speed and growth rate: (a) Front migration rate (heatmap) as a function of run speed (|𝑣_*r*_|) and maximum growth rate (*k_g_*) simulated using the reaction-diffusion model discussed in the text with parameters appropriate for rich medium conditions (Table 11). Model parameters were estimated using the method developed by Croze *et al* (see supplementary file 1). Gray square shows the run speed and growth rates measured for the founding strain in rich medium (see Figure 3). Standard error in |𝑣_*r*_| is smaller than the size ofthe marker; error bar in *k_g_* is the standard deviation across three replicate measurements. (b) Identical to panel (a) except for minimal medium. The abrupt change in migration rate around *k_g_* =0.2h^−1^ corresponds to a transition from diffusion dominated front migration to a traveling wave (see supplementary file 1). The founding strain’s phenotype is shown as a red circle, error bars are constructed identically to those in (a). Figure 2 - figure supplement 1: Reaction-diffusion model recapitulates qualitative features of colony expansion. Figure 2 - figure supplement 2: Comparison of front profiles from simulation and experiment, Figure 2 - figure supplement 3: Simulation of migration rate versus tumble frequency.

To understand how changes in motility and growth could contribute to the evolution of migration, we studied how the migration rate (*s*) varied with the parameters of our model through numerical simulation (supplementary file 1). We found that increases run speed (|𝑣_r_|) and growth rate (*k_g_*) had the largest impact on *s* (Figure 2). Consistent with previous reports, our model indicates that only small gains in migration can be achieved through increases in tumble frequency (*Wolfe and Berg, 1989*) (~10%, Figure 2 - figure supplement 3).

Figure 2 shows how the front migration rate (heatmap) varies with run speed and growth rate for both nutrient conditions studied in Figure 1. Our model predicts that the fastest migrating strain should be the one that increases both its run speed and growth rate relative to the founder. Therefore, in the absence of any constraints on accessible phenotypes, we expect both run speed and growth rate to increase with selection.

### A trade-off constrains the evolution of faster migration

To test the predictions of the reaction-diffusion model, we experimentally interrogated how the motility and growth phenotypes of our populations evolved over the course of selection. We performed single-cell tracking experiments using a microfluidic method similar to one described previously (*Jordan et al., 2013*). This method permitted us to acquire 5 minute swimming trajectories from hundreds of individuals from strains isolated prior to selection (founder) and after 5,10 and 15 rounds of selection in rich media (replicate 1, Figure 1(c)) and for the founder and strains isolated after 5 and 10 rounds of selection in minimal media (replicate 1, Figure 1(e)). For tracking, cells were grown in the medium in which they were selected. This technique permitted us to capture more than 280 000 run-tumble events from approximately 1500 individuals.

We identified run and tumble events for all individuals (*Berg and Brown, 1972; Tauteetal., 2015*) (Methods). Figure 3(a-b) shows that run durations declined over the course of selection in both rich and minimal media. We show the complementary cumulative distribution function (*c*(𝜏_*r*_)) of run durations (𝜏_*r*_) aggregated across all run events detected for the founding or evolved strains 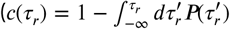 where 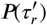 is the distribution of run durations). *c*(𝜏_*r*_) quantifies the fraction of all runs longer than a time 𝜏_*r*_. These distributions show that the evolved strains exhibited a reduction in the probability of executing long runs. This reduction in run duration (increase in tumble frequency) is expected since previous studies showed that mutants with longer run durations have slower migration rates through agar (*Wolfe and Berg, 1989*). We observed opposite trends for tumble duration, with decreasing tumble duration in rich medium and increasing duration in minimal medium (Figure 3 - figure supplement 2).

**Figure 3.**
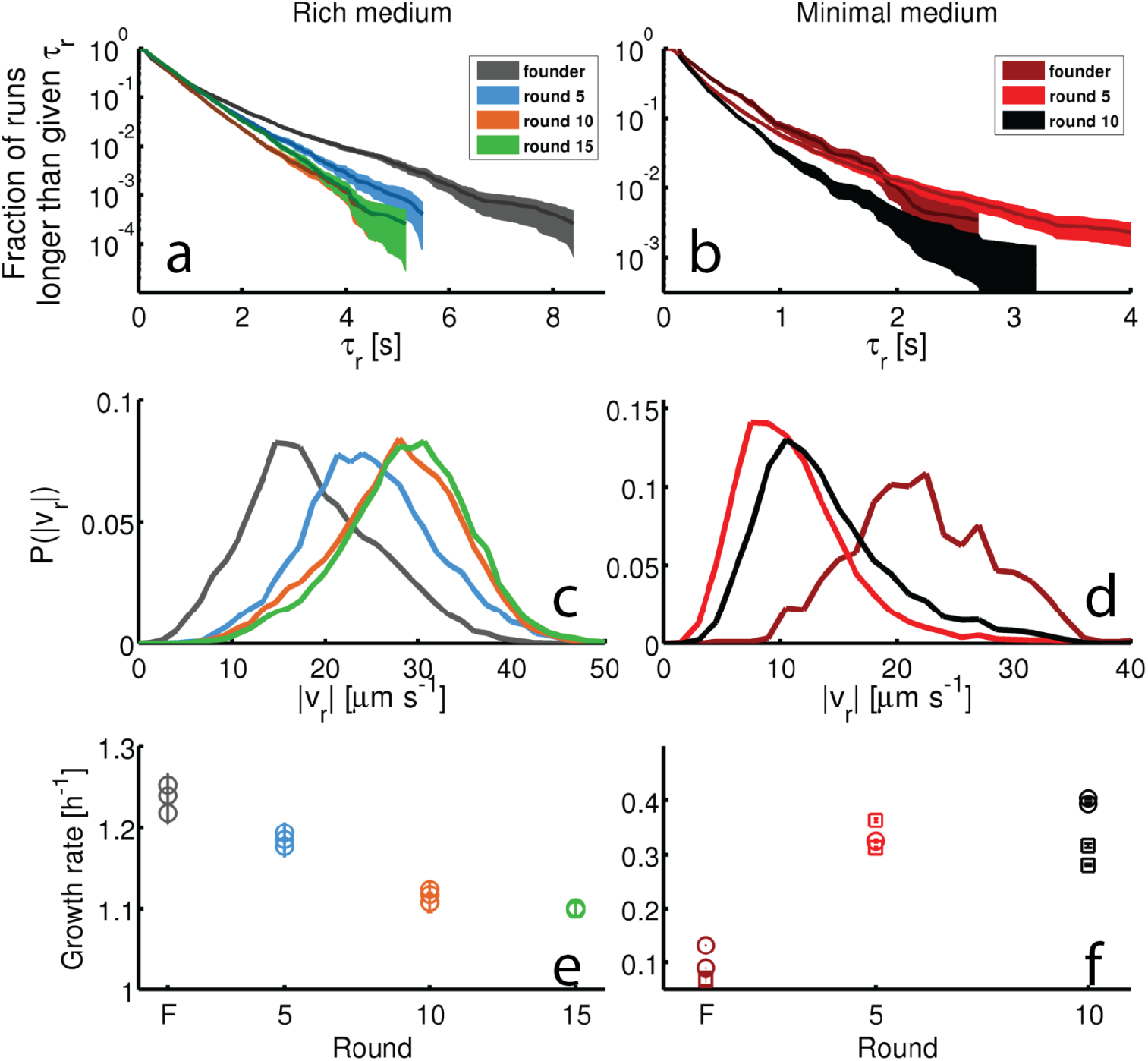
Dynamics of phenotypic evolution in rich and minimal media: (a-d) Show single-cell swimming phenotypes (run duration (*𝜏_r_*) and run speed (|𝑣_*r*_|)) (see Methods). Tracking was performed for founding strain (140 cells, 19 597 run events), strains isolated after 5 (79 cells, 12217 run events), 10 (97 cells,18505 run events) and 15 (96 cells, 15928 run events) rounds in rich media and in minimal media for the founding strain (20 cells,1752 run events), round 5 (45 cells, 9724 run events) and round 10 (29 cells, 5384 run events). (a) Shows the fraction of runs longer than a given *𝜏_r_* for strains evolved in rich media (95 % confidence intervals from bootstrapping). The mean and standard deviation in run duration for founder is 0.66±0.78 s, for round 5: 0.63±0.61 s, for round 10: 0.58±0.50s and for round 15: 0.65±0.57s. Round 5, 10 and 15 strains exhibit shorter average run durations than founder (*p* <0.05). (b) Shows the same distribution for strains in minimal medium with founder exhibiting average run duration 0.47±0.45 s, round 5: 0.4±0.47 s and round 10: 0.34±0.30s. Rounds 5 and 10 exhibit shorter average run durations than founder (*p* <10^−8^). (c) Shows run speed distributions for strains evolved in rich medium, legend in (a) applies. The average ± standard deviation run speeds are, for founder: 18.7±7.1 μms^−1^, round 5: 24.9±7.1 μms^−1^, round 10: 27.6±7μms^−1^, and for round 15: 28.7±6μms^−1^. Average run speeds for rounds 5, 10 and 15 are greater than founder (*p* <10^−5^). (d) Shows the same distributions for strains evolved in minimal medium, average run speed for founder: 22.2±6.1 μms^−1^, for round 5: 11.2±5μms^−1^ and for round 10: 13.9±5.9 μms^−1^. Both rounds 5 and 10 exhibit slower average run speeds than founder (*p* <10^−5^). Legend in (b) applies. (e-f) Show growth rates in well mixed liquid culture for all strains studied in panels (a-d) in the medium in which the strains were selected. (e) Shows triplicate measurements from each of the four strains isolated in rich medium. Rounds 5, 10 and 15 exhibit slower growth than founder (*p*<0.01). (f) Shows growth rates for strains isolated from minimal medium selection experiment. Four replicate measurements were made for founder and round 10 and three replicate measurements for round 5. Squares and circles demarcate measurements made on separate days. Rounds 5 and 10 have higher growth rates than founder (*p* <10^−5^). Figure 3 - figure supplement 1: Microfluidic device and single-cell swimming trajectory, Figure 3 - figure supplement 2: Tumble durations and run lengths for evolved strains, Figure 3 - figure supplement 3: Reproducibility of the evolved phenotype, Figure 3 - figure supplement 4: Swimming statistics as a function of culture density.

Figure 3(c-d) show the probability distributions of run speeds for founding and selected strains in both nutrient conditions. In rich medium we observed a nearly 50% increase in the run speed (|𝑣_*r*_|) between founder and rounds 10 to 15. Tracking strains isolated after 15 rounds from independent selection experiments (replicates 3 and 4, Figure 1(c)) showed that this increase in run speed was reproducible across independent evolution experiments (Figure 3 - figure supplement 3). Finally, to check that the phenotype we observed after 15 rounds of selection in rich medium was distinct from standard laboratory strains used in chemotaxis studies we tracked RP437 and found that its swimming speed was slower than the round 15 strain (Figure 1 - figure supplement 4).

Surprisingly, when we performed single-cell tracking for strains evolved in minimal media we observed the opposite trend. In these conditions we observed a 50 % reduction in run speed (Figure 3(d)). Again, we found that this result was reproducible across independently evolved strains (Figure 3 - figure supplement 3).

The reduction in run duration observed for the strain isolated after 10 rounds of selection in minimal medium was not reproducible, with an independently evolved strain exhibiting very long runs (Figure 3 - figure supplement 3). The strain where we observed long runs after 10 rounds of selection (replicate 2, Figure 3(e)) also exhibited a slower migration rate than the strain isolated from replicate 1, and the long run durations may be responsible for this difference.

We then measured the growth rates in well mixed liquid of founding and evolved strains from both selection conditions in Figure 1 (Methods). We observed a decline of about 10% in the maximum growth rate with selection in rich medium and a three-fold increase in the maximum growth rate after 10 rounds of selection in minimal medium (Figure 3(e,f)). We found that these changes in growth rate are reproducible across independently evolved strains in both environmental conditions (Figure 3(e,f), Figure 3 - figure supplement 3).

Since motility is known to depend on the growth history of the population (*Staropoli and Alon, 2000*), we checked whether the conclusions made above remained valid when we tracked cells over a range of optical densities during population growth. We performed these measurements for the founding strain in both rich and minimal media, and for a round 15 strain in rich medium and a round 10 strain in minimal medium and confirmed that the differences in |𝑣_*r*_| and 𝜏_*r*_ were retained across phases of growth (Figure 3 - figure supplement 4).

Combining growth rate measurements with single-cell motility measurements allowed us to predict the front migration rate for strains in rich and minimal medium using the reaction-diffusion model described above. We found that the model qualitatively recapitulated the increase in front migration rate that we observed experimentally (Tables 12 and 13, Figure 4 - figure supplement 1).

**Table 12.** Reaction-diffusion model parameters estimated from measurements of tumble frequency (*α*) and run speed (|𝑣_*r*_|) for rich medium evolved strains in *C* =0.3 % agar.

| Evolution of population level migration parameters |  |  |  |  |
| --- | --- | --- | --- | --- |
| strain | $\alpha_0 [\text{s}^{-1}]$ | $v_r [\mu\text{m s}^{-1}]$ | $D_b [\text{cm}^2 \text{h}^{-1}]$ | $k_0 [\text{cm}^2 \text{h}^{-1}]$ |
| founder | 1.45 | 18.7 | 0.02 | 0.65 |
| round 5 | 1.56 | 24.9 | 0.027 | 0.90 |
| round 10 | 1.72 | 27.6 | 0.029 | 1.04 |
| round 15 | 1.54 | 28.7 | 0.031 | 1.04 |

**Table 13.** Reaction-diffusion model parameters estimated from measurements of tumble frequency (*α*) and run speed (|𝑣_*r*_|) for minimal medium evolved strains in *C* =0.3 % agar.

| Evolution of population level migration parameters |  |  |  |  |
| --- | --- | --- | --- | --- |
| strain | $\alpha_0$ [$s^{-1}$] | $v_r$ [$\mu m s^{-1}$] | $D_b$ [$cm^2 h^{-1}$] | $k_0$ [$cm^2 h^{-1}$] |
| founder | 2.125 | 22.2 | 0.021 | 0.66 |
| round 5 | 2.47 | 11.2 | 0.0099 | 0.35 |
| round 10 | 2.88 | 13.2 | 0.011 | 0.44 |

**Figure 4.**
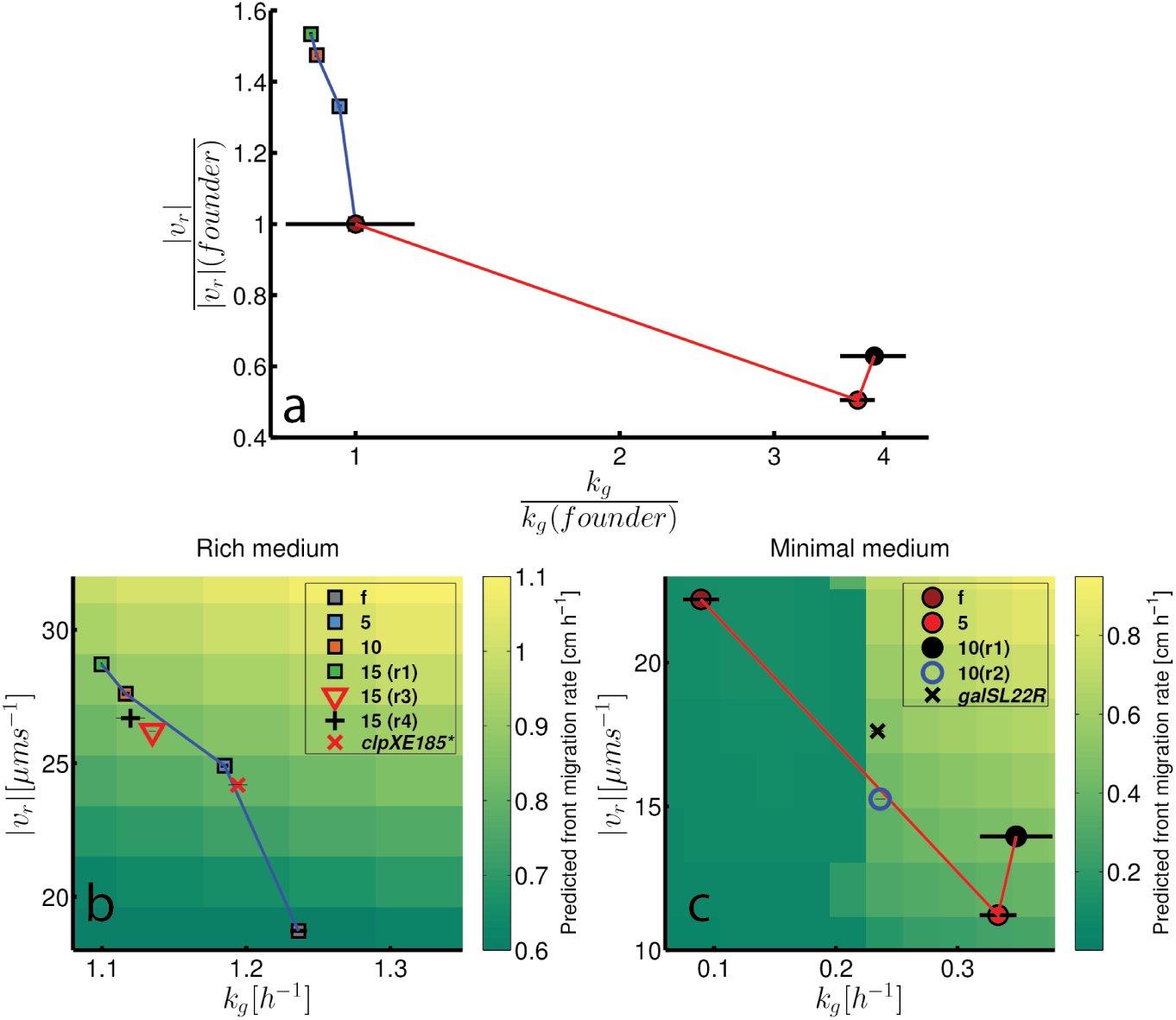
Trade-off between growth rate and run speed constrains evolution of faster migration: (a)Run speeds and growth rates for strains evolved in rich medium (squares) and minimal medium (circles) normalized to the run speed and growth rates of the founding strain in each environment. Standard error in run speeds are smaller than the markers. The large error bar at the point (1,1) corresponds to the large error in the growth rate measurement of the founding strain in minimal medium. (b) Reproduces the data from (a) for evolution in rich medium overlaid on a heatmap of the prediction for front migration rate from the reaction-diffusion model (Figure 2). Growth rates and run speeds are not normalized. Phenotypes for strains from Figure 3 are shown along with two strains from independently evolved strains (replicates 3 (15(r3)) and 4 (15(r3)), Figure 1). In addition, the red “x” marks the phenotype for the mutation (clpXE185*) in the founding strain background (see Figure 5). (c) Shows run speeds and growth rates for strains evolved in minimal medium overlaid on the predicted from migration rate from the reaction-diffusion model. Growth rate and run speed for an independently evolved round 10 strain is shown (10(r2)) as well as the phenotype for the galSL22R mutation in the founder background (black “x”). Predicted front migration rates assume no change in run duration. Figure 4 - figure supplement 1: Predicted migration rates for evolved strains, Figure 4 - figure supplement 2: Swimming statistics, growth rates and migration rates for mutants.

We conclude that there is a trade-off between run speed and growth rate in *E. coli* which constrains the evolution of faster migration through low viscosity agar. Figure 4 summarizes this trade-off for both rich medium and minimal medium selection experiments. Figure 4(a) shows the changes in run speed and growth rate normalized to the values for the founding strain in each condition. The curve in Figure 4(a) constitutes a Pareto frontier in the phenotypic space of run speed and growth rate. In Figure 4(b-c) we show the measured growth rates and swimming speeds for all strains presented in Figure 3 overlaid on the predicted migration rates from our reaction-diffusion model.

### Parallel genomic evolution drives a trade-off through antagonistic pleiotropy

To investigate the mechanism of the phenotypic evolution and tradeoff we observed, we performed whole genome sequencing of populations for the founding strain as well as strains isolated after rounds 5, 10 and 15 in rich medium for four of five selection experiments and rounds 5 and 10 in minimal medium for four of five selection experiments (Methods). Figure 5 shows *de novo* mutations observed in each strain sequenced. Since we sequenced populations, we report the frequency of each mutation observed (see legend, Figure 5(a), middle panel).

**Figure 5.**
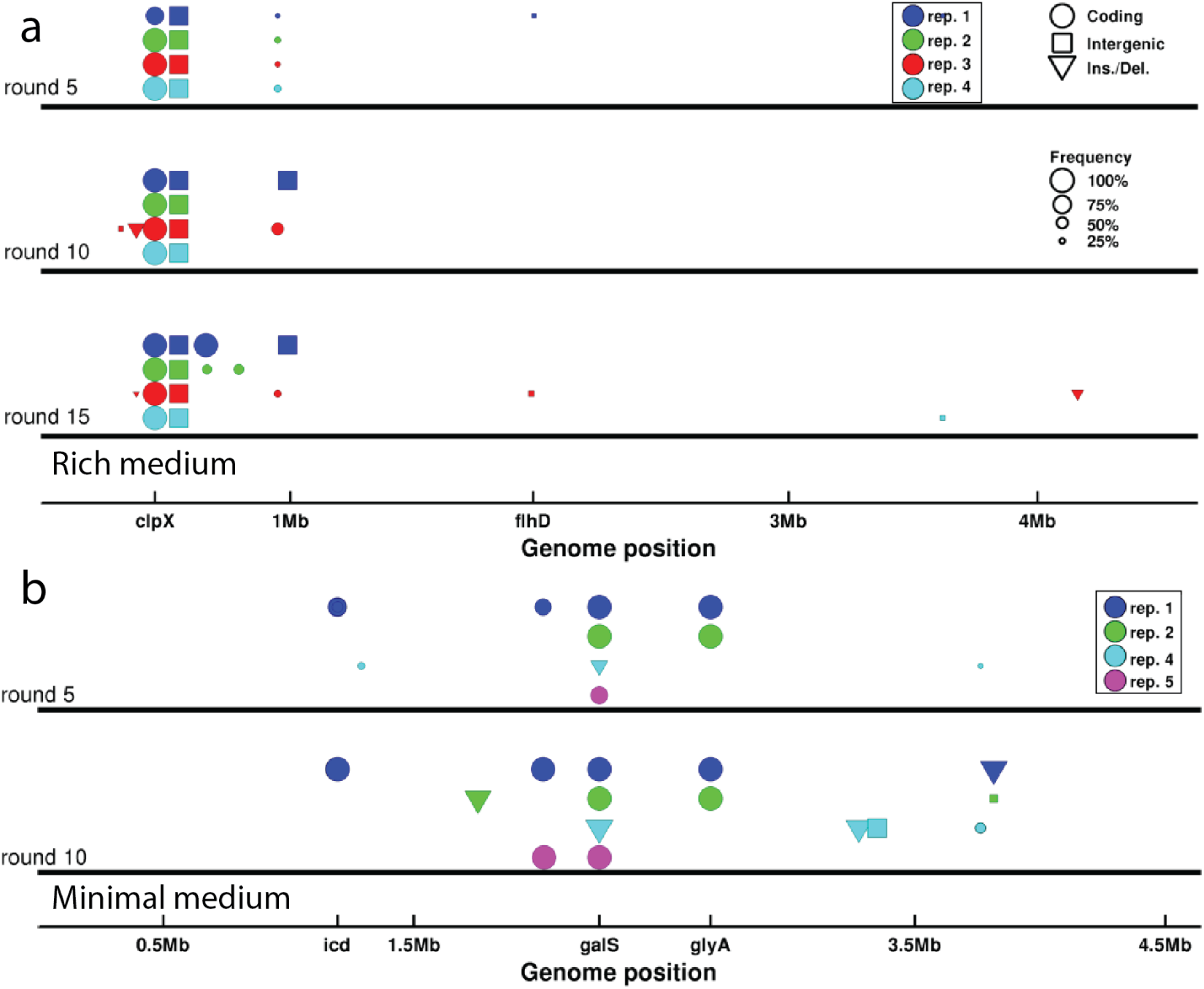
Genomic evolution: (a) *De novo* mutations observed in strains isolated after 5, 10 and 15 rounds of selection in rich medium. Abscissa denotes position along the genome. Colors of the markers indicate independently evolved replicates and correspond to traces in Figure 1(c). Circles denote single nucleotide polymorphisms (SNP) in coding regions, squares denote intergenic SNPs, and triangles denote larger insertions or deletions. The size of the marker is proportional to the frequency of the mutation in the population. Only mutations with a frequency above 0.2 in the population are shown. Genes of interest are shown. The operons coding for motility and chemotaxis are near *flhD*. (b) Identical to (a) but shows *de novo* mutations for strains evolved in minimal medium. The marker near *icd* corresponds to multiple SNPs in close proximity to each other. See Tables 1 to 4 and 6 to 9 for a list of all mutations observed and details ofthe sequencing.

In the rich medium experiment we observed parallel evolution across replicate selection experiments, with a mutation in *clpX* (E185*) and an intergenic single base pair deletion both rising to fixation within approximately 5 rounds of selection. In this condition we observed transient mutations in genes regulating chemotaxis or motility (near *flhD*, Figure 5(a)) in two of four replicates.

A previous study showed that mutations in *clpX* alter *flhDC* expression and motility (*Girgis et al., 2007*). We therefore focused attention on the mutation in *clpX*, which converted position 185 from glutamic acid to a stop codon in the 424 residue *ClpX* protein. *ClpX* is the specificity subunit of the *ClpX-ClpP* serine protease. *ClpX* forms a homohexamer that consumes ATP to unfold and translocate target proteins to the *ClpP* peptidase (*Baker and Sauer, 2012*). The *ClpXP* protease has many targets in the cell including *FlhDC*, the master regulator of flagellar biosynthesis (*Tomoyasu et al., 2003*). We found that this mutation in *clpX* was at high abundance (>70%) in all populations after 5 rounds of selection and fixed by round 10 in all four replicates (Figure 5).

To determine the phenotypic effects of *clpX*E185*, we used scarless recombineering to reconstruct this mutation in founding strain genetic background (*Kuhlman and Cox, 2010*) (Methods). We then performed migration rate, single-cell tracking and growth rate measurements on this strain (Figure 4 - figure supplement 2). We observed a statistically significant increase in migration speed for the *clpX*E185* mutant (0.39±0.01 cmh^−1^, mean and standard error) relative to founder(0.30±0.01 cmh^−1^, *p* =0.002, Figure 4 - figure supplement 2). We also found that *clpX*E185* resulted in a statistically significant increase in run speed relative to founder (24.18 μm^−1^ compared to 18.7μm^−1^, *p* <10^−10^). Finally, in well mixed batch culture in rich medium the *clpX*E185* mutant exhibited a maximum growth rate *k_g_* = 1.19±0.009h^−1^ (standard error for triplicate measurements) with founder exhibiting a maximum growth rate of 1.23±0.01 h^−1^ (p = 0.0174). Knocking out *clpX* from founder resulted in very slow front migration (*s* =0.0036±0.001 cmh^−1^), suggesting that the stop codon mutation we observe has a more subtle effect on the enzyme’s function than a simple loss of function. Finally, we reconstructed the intergenic single base pair deletion which fixed in all four replicate selection experiments but observed no phenotypic effects of this mutation when placed in the founder or *clpX*E185* background (Figure 4 - figure supplement 2). These results suggest that this intergenic mutation is neutral.

We conclude that the *clpX* mutation observed in all four replicate experiments drives faster front migration through increasing run speed, despite decreasing growth rate. Since the mutant’s phenotype lies on the Pareto frontier in Figure 4(b) we conclude that the trade-off between growth rate and swimming speed is driven by antagonistic pleiotropy (*Cooper and Lenski, 2000*).

Figure 5(b) shows the mutations observed in rounds 5 and 10 for four of five replicate selection experiments in minimal medium. In all experiments we observed mutations in the transcriptional regulator *galS* which fixed in just 5 rounds. In one of four experiments we observed a mutation in the gene encoding the motor protein *FliG*, otherwise the observed mutations appear to be metabolic in nature. In minimal medium we also observed a substantial number of synonymous mutations rising to fixation (see tables 6 to 9). The role of these synonomous mutations is not known, but may be due to tRNA pool matching (*Stoletzki and Eyre-Walker, 2007*).

To understand how these mutations drive phenotypic evolution, we focused on the galSL22R mutation. *galS* encodes the transcriptional repressor of the *gal* regulon. The coding mutation we observe occurs in the highly conserved N-terminal helix-turn-helix DNA binding region of this protein, we therefore expect that this mutation alters the expression of the *gal* regulon (*Weickert and Adhya, 1992*). To assay the phenotypic effects of this mutation, we reconstructed it in the genetic background of the founder.

The migration rate of the galSL22R mutant showed a statistically significant increase relative to founder (*s* =0.039±0.001 cmh^−1^ for galSL22R and 0.0163±0.0038cmh^−1^ for founder, *p* <10^−3^). We found that the growth rate of the mutant was approximately 2.5-fold larger than founder in minimal medium (0.23±0.005 h^−1^ for galSL22R and 0.089±0.03 h^−1^ for founder, *p* =4 × 10^−4^). Further, this mutation reduced the mean swimming speed relative to founder by approximately 20% (Figure 4(c),(Figure 4 - figure supplement 2). However, when we knock out the *galS* gene from founder we do not observe a significant increase in the migration rate (Δ*galS s* =0.0165±0.002 cm h^−1^, *p* = 0.92).

Therefore, as shown in Figure 4(c), we conclude that *galS*L22R alone is capable of driving the population in the opposite direction along the Pareto frontier observed in Figure 4(a). As with the rich medium condition, this trade-off is governed by antagonistic pleiotropy.

### Genetic covariance determines direction of phenotypic evolution

To understand why we observe divergent phenotypic trajectories in the rich and minimal medium conditions (Figure 4(a)), we studied a simple model of the evolution of correlated traits (*Lande, 1979; Mezey and Houle, 2005*). We consider a vector of the two phenotypes of interest, run speed and maximum growth rate, normalized to the values ofthe founder (*Hansen and Houle, 2008*), 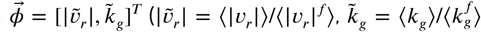, where 〈〉 denotes an average across the population). The model describes the evolution ofthe mean phenotype 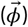 under selection by

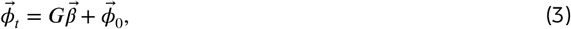

where ***G***, the genetic covariance matrix, describes the genetically driven phenotypic covariation in the population, which is assumed to be normally distributed 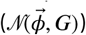. 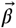 is the selection gradient which captures the change in migration rate with respect to phenotype since we are selecting for faster migration. The matrix ***G*** is given by

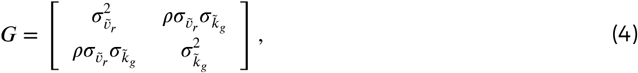

where 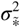 describes the (fractional) variance in the phenotype due to genetic variation and *ρ* captures the correlation between the two traits. In our experiment we do not have a direct measurement of ***G***. However, we do observe how 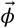 changes over the course of selection, our data suggest that *ρ* < 0 and our reaction-diffusion model permits us to estimate how migration rate depends on the two traits of interest. In particular, 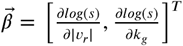. We approximate 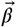 in both rich and minimal media by fitting a plane to the heatmap shown in Figure 2(a-b) (supplementary file 1). The resulting selection gradient is shown in Figure 6 - figure supplement1 for both conditions. Using this formalism, we asked what values of 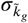 and 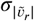 would result in the directions of phenotypic evolution we observed experimentally in rich and minimal media.

**Figure 6.**
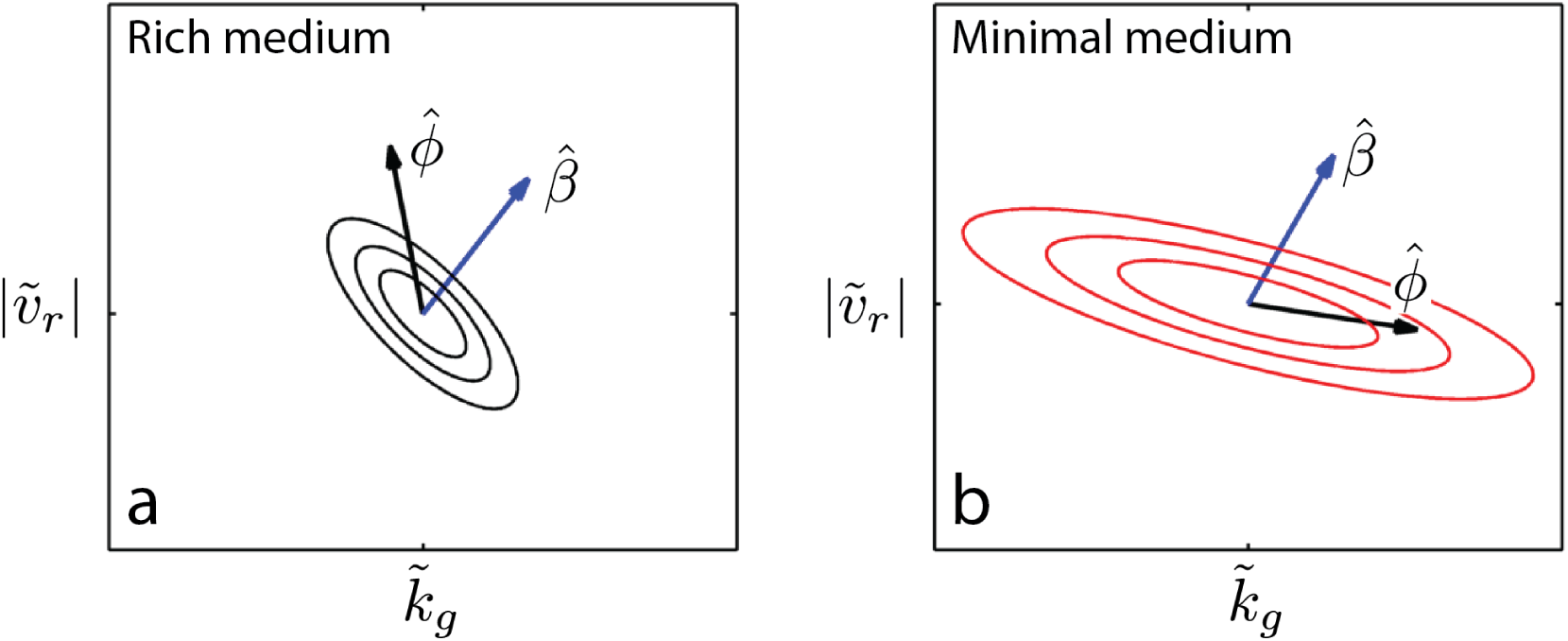
Evolution of correlated traits: The evolutionary model describes the change in phenotype relative to the founder 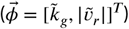 under selection described by 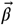. Panels show unit vectors in the direction of observed phenotypic evolution 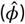 and the direction of selection inferred from the reaction-diffusion model 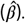 Ellipses show quartiles for a normal distribution of phenotypes with covariance matrix *G* that is consistent with 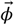 and 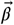. In both panels we set the correlation coefficient between 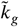 and 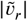 is *𝜌* = −0.75 but our conclusions hold for *𝜌* < −0.1. In rich medium (a) 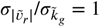 and in minimal medium 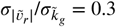. In rich medium *β̂_RM_* = [0.61,0.78] and in minimal medium *β̂_MM_* = [0.5,0.87]. Figure 6 - figure supplement 1: Determining 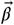 from reaction-diffusion model, Figure 6 - figure supplement 2: Direction of phenotypic evolution with 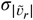 and 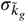, Figure 6 - figure supplement 3: Stochastic simulations of selection in minimal medium.

We found that the direction of phenotypic evolution in rich medium agreed well with our experimental observations so long as 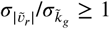 for *ρ* < −0.1. This implies that our observed phenotypic evolution is consistent with a genetic variance in run speed that is no smaller than the genetic variance in growth rate (Figure 6 - figure supplement 2). In contrast, in minimal medium the model predicts the direction of observed phenotypic evolution only if 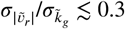 for *ρ* < −0.1. This result indicates that our observed phenotypic evolution is consistent with at least three-fold larger propensity for mutations to alter growth rate compared to run speed in minimal medium (Figure 6 - figure supplement 2).

This suggests that the capacity of mutations to alter run speed or growth rate relative to founder depends on the nutrient conditions and that changes in this capacity qualitatively alter the direction of evolution along the Pareto frontier. This result captures the intuition that mutations that can increase growth rate in rich medium are few while in minimal medium the propensity for mutations increasing growth rate is substantially larger. The model presented here relies on a linear approximation to 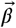 which is a good assumption for rich medium but not for minimal medium, where the dependence of *s* on |𝑣_*r*_| and *k_g_* is strongly nonlinear. Using simulations of the evolutionary process described by equation 3, we relaxed the assumption of linearity in the selection coefficient and found that our qualitative conclusions were not altered (Figure 6 - figure supplement 3).

We note that the structure of ***G*** inferred above reflects the capacity for mutations to change phenotypes at the outset of the experiment. As evolution proceeds in rich medium, we observe a saturation in both run speed and growth rate (Figure 4(c,e)), suggesting that further variation along the Pareto frontier is constrained, either genetically or through biophysical constraints on swimming speed. Similarly, in minimal medium, saturation in the growth rate occurs after 5 rounds of selection, suggesting that mutations to further improve growth rate are either not available or fundamental constraints on growth inhibit further increases (*Scott et al., 2010*).

## Discussion

The most striking observation of our study is the divergent trajectories of phenotypic evolution along a Pareto frontier (Figure 4(a)). This observation shows that the evolution of faster migration results in environmentally dependent phenotypic outcomes. This result has important implications for interpreting phenotypic variation in natural populations.

When trade-offs are observed in wild populations it is sometimes proposed that phenotypes at the extrema of a Pareto frontier reflect the outcome of selection for a specific task (*Shoval et al., 2012*). Our study shows that when selection pressures place demands on multiple traits simultaneously, evolution along the frontier can reflect differing genetic capacity for adaptation of each phenotype rather than simply the fitness benefit of improving each trait. This result suggests a cautious approach to interpreting phenotypes in Nature where selection pressures and mechanisms constraining phenotypes are often not known (*Gould and Lewontin, 1979*).

Our results point to the potential predictive power of determining the directions in phenotype space in which genetic variation can most readily change phenotypes - so called, 'genetic lines of least resistance’ (*Schluter, 1996*). These directions may be related to genetic regulatory architecture. The mutations we observe in both rich and minimal media alter negative regulators (a protease in the case of *clpX* and a transcriptional repressor in the case of *galS*). This supports the hypothesis that microevolution is dominated by the disruption of negative regulation (*Lind et al., 2015*) and suggests that the direction of phenotypic evolution can be predicted by determining where negative regulatory elements reside in genetic and proteomic networks. The mutations we examined appear to be more subtle than simple loss of function, since knockout mutants for both *clpX* and *galS* do not exhibit fast migration, therefore a detailed understanding of how mutations disrupt negative regulation will be essential.

The trade-off presented here has been observed previously in *E.coli* (*Yi and Dean, 2016*) and Pseudomonas aeruginosa (*Deforet et al., 2014; van Ditmarsch et al., 2013*). Yi and Dean selected *E. coli* alternately for growth and chemotaxis and observed a trade-off between growth rate and swimming speed which was circumvented by phenotypic plasticity. We observe no evolution beyond the Pareto frontier in our study, possibly because our conditions simultaneously select for growth and motility rather than alternating between selection pressures. This suggests that evolutionarily persistent trade-offs may reflect selection pressures that occur simultaneously in Nature. Both van Ditmarsch *et al.* (*van Ditmarsch et al., 2013*) and Yi and Dean (*Yi and Dean, 2016*) observe mutations that alter regulation of motility and chemotaxis genes. Interestingly, none of the mutations observed in our experiment were found by Yi and Dean, despite evolution along similar Pareto frontiers. This suggests that determining the allowed directions of phenotypic variation may be a more powerful approach to predicting evolution than cataloging mutations alone.

The mechanism of the trade-off between growth rate and swimming speed has, to our knowledge, not been determined. However, over-expression of motility operons could drive the reductions in growth rate we observe in rich medium. Subsequent increases in speed could then arise passively from reductions in cell size which reduce hydrodynamic drag (*Taheri-Araghietal., 2015*). Similarly, increases in growth rate in minimal medium should increase cell size and hydrodynamic drag. Using the data of Taheri-Araghi *et. al* (*Taheri-Araghi et al., 2015*) we estimated changes in cell size due to measured changes in growth rate for populations evolved in rich and minimal medium. We could not account for the large change in swimming speed we observe through growth rate mediated changes in cell size alone (supplementary file 1). Since we have not measured cell size directly, we cannot conclusively rule out this mechanism. To definitively characterize the mechanism of this trade-off will require measurements of cell size, gene expression, flagellar length and proton motive force.

Our study shows how evolutionary dynamics are defined by the complex interplay between genetic architecture, phenotypic constraints and the environment. Our hope is that a general approach to predicting evolution can emerge from a more complete understanding of this interplay.

## Methods

### Motility Selection

#### Rich medium

10μL of motile *E. coli* (strain MG1655-motile, Coli Genetic Stock Center (CGSC) #8237) from an overnight LB culture was injected at the center of a 0.3 % w/v agar 15 cm diameter plate containing LB. Images were acquired every minute via webcams (Logitech HD Pro Webcam C920, Logitech, Lausanne, Switzerland) in a dark box using pulsed illumination provided by an LED light strip (part number: COM-12021, Sparkfun Inc, Nilwot, CO). Only the Red and Green LEDs of the RGB strip were used in order to avoid blue light response known to occur in *E. coli* (*Taylor and Koshland, 1975*). After 12 hours, 50 μL of cells was removed from each of eight points around the outermost ring. This sample was briefly vortexed and 10μL (≈ 10^6^ cells) was injected into a fresh plate from the same batch. Remaining bacteria were preserved at −80°C in 25 % glycerol. Selection was performed by repeating this sampling and growth over 15 rounds. Automated image processing then yielded quantitative data about front geometry and kinetics. All experiments were performed at 30°C in an environmental chamber (Darwin Chambers, St. Louis, MO). Plates were thermalized for 12 hours in the environmental chamber before use.

To estimate the number of generations that occur during each round of selection, we inoculated an agar plate from a culture of the founding strain and sampled the migrating front as in the selection experiment. We then measured the cell density of the sampled population by serial dilution and plating. We inoculated a fresh plate (with 10^6^ cells) and permitted the colony to expand for 12 hours. To measure the total population on the plate after growth, we mixed the entire contents of the plate in a beaker and measured the density by serial dilution and plating again. From this we extracted an estimate of the number of generations that occurred. The range reflects the large errors due to serial dilution and plating.

#### Minimal medium

Selection experiment was performed identically to rich medium experiment with the following modifications. Plates were made with M63 0.18mM galactose. Cultures used to initiate selection were grown in M63 30mM galactose for 24 to 48 hours prior to initiating selection. During each 48 hour round of migration and imaging, plates were housed in a plexiglass box with a beaker of water to prevent evaporative losses from the plate. Images were acquired every 2 minutes. We estimated the number of generations per round as described above. Reliable plate counts were only obtained for plates of round 10 strains where we estimate 10 generations per round. We therefore take this as an upper bound and conclude that the 10 round selection experiment includes <100 generations. Plates were thermalized for 24 hours before use.

The Δ*cheA-Z* mutant was constructed via P1 transduction from a strain provided by the group of Chris Rao and the mutation was confirmed by PCR. This mutant lacks the receptors *tar* and *tap* and the chemotaxis genes *cheAWRBYZ*.

We selected the motile MG1655 wild-type strain for these experiments rather than the more commonly used RP437 strain since the latter is auxotrophic for several amino acids. Minimal medium experiments were therefore performed without additional amino acids which could confound results.

### Image analysis

Webcam acquired images of migrating fronts were analyzed by custom written software (Matlab, Mathworks, Natick, MA). A background image was constructed by median projecting 6 images from the beginning of the acquisition before significant growth had occurred. This image was subtracted from all subsequent images prior to further analysis. The location of the center of the colony was determined by first finding the edges of the colony using a Canny edge detection algorithm. A circular Hough transform (*Hough, 1959*) was applied to the resulting binary image to locate the center. In rich medium, where signal to background was > 10, radial profiles of image intensity were measured from this center location and were not averaged azimuthally due to small departures from circularity in the colony. The location of the front was determined by finding the outermost peak in radial intensity profiles. Migration rate was determined by linear regression on the front location in time. Imaging was calibrated by imaging a test target to determine the number of pixels per centimeter. The results of the calibration did not depend on the location of the test target in the field of view. In minimal medium, where the signal to background is reduced due to low cell densities, background subtraction was employed as described above but radial density profiles were not always reliable for locating the front. Instead, a circular Hough transform was applied to each image to locate the front at each point in time.

### Single-cell tracking

Single-layer microfluidic devices were constructed from polydimethyl-siloxane (PDMS) using standard soft-lithography techniques, (*Quake and Scherer, 2000*) following a design similar to one used previously (*Jordan et al., 2013*), and were bonded to coverslips by oxygen plasma treatment (Harrick plasma bonder, Harrick Plasma, Ithaca, NY). Bonded devices formed a circular chamber of diameter 200 μm and depth 10μm (Figure 3 - figure supplement 1). Devices were soaked in the medium used for tracking (LB for rich medium strains, M63 0.18mM galactose for minimal medium strains) with 1 % Bovine Serum Albumin (BSA) for at least 1 hour before cells were loaded. Bacteria were inoculated directly from frozen stocks into medium containing 0.1 % BSA in a custom continuous culture device. BSA was necessary to minimize cells adhering to the glass cover slip. For rich medium tracking experiments cells grew to a target optical density and the continuous culture device was run as a turbidostat. In minimal medium experiments the device was run as a chemostat at an optical density of ~0.15. The culture was stirred by a magnetic stir bar at 775 RPM and the temperature was maintained at 29.75°C by feedback.

To perform single-cell tracking, cells were sampled from the continuous-culture device and diluted appropriately (to trap one cell in the chamber at a time) before being pumped into the the microfluidic chamber. Video was acquired at 30 frames per second with a Point Grey model FL3-U3-32S2M-CS camera (Point Grey, Richmond, Canada) and a bright-field microscope (Omano OM900-T inverted) at 20× magnification. Movies were recorded for 5 minutes before a new cell was loaded into the chamber. Two microscopes were operated in parallel. The stock microscope light source was replaced by a high-brightness white LED (07040-PW740-L, LED Supply, Randolf, VT) to avoid 60Hz flickering that was observed with the stock halogen light source. All experiments were performed in an environmental chamber maintained at 30°C.

Movies were segmented and tracked with custom written Matlab routines described previously (*Jordan et al., 2013; Jaqaman et al., 2008*). At times when two individuals are present in the chamber, ambiguous crossing events can lead to loss of individual identities. All crossing events were inspected manually to prevent this. To identify runs and tumbles, we utilized a method based on reference (*Taute et al., 2015*) which was modified from the approach used by Berg and Brown (*Berg and Brown, 1972*). Briefly, for each cell the segmentation routine results in a matrix of spatial locations 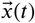. We compute the velocity by the method of central differences resulting in 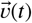 from which we compute an angular velocity between adjacent velocity vectors (*𝜔*(*t*)). We then define *α*, a threshold on *𝜔*. Tumbles are initiated if *𝜔*(*t*) & *𝜔*(*t* + 1) > *α* or if *𝜔*(*t*) > *α* and the angle defined between the vectors 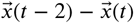 and 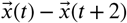 is greater than *α*. The latter condition detects tumbles that occur on the timescale of the imaging (0.033 s). Runs are initiated only when *𝜔*(*t*) & *𝜔*(*t* + 1) & *𝜔*(*t* + 2) < *α*. As a result, tumbles can be instantaneous and runs are a minimum of four frames. *α* was determined dynamically for each individual by initializing *α*_0_ and then detecting all runs for a cell. A new *α_i_* = *c* × median(*𝜔_runs_*) was computed with *c* a constant and *𝜔_runs_* is the angular velocity during runs. The process was iterated ten times but typically converged to a final *α_f_* in less than five iterations. *c* = 5 was determined by visual inspection of resulting classified trajectories. Approximately 1 % of cells exhibited sustained tumbling and had average tumble durations greater than 0.4 s and were excluded from further analysis. We only considered run events that were not interrupted by interactions with the circular boundary of the chamber.

Due to interactions with the chamber floor and ceiling (boundaries perpendicular to the optical axis) we intermittently observed cells circling. We developed a method to detect this behavior automatically and found that our results are unchanged when we consider individuals that are not interacting the the chamber boundaries (supplementary file 1).

### Whole genome sequencing and analysis

Whole genome sequencing was performed using the Illumina platform with slight variations between four independent runs. For all sequencing, cultures were grown by inoculating fresh medium from frozen stocks isolated during the course of selection and growing to saturation at 30°C. For sequencing of rich medium strains from replicate 1, DNA was extracted and purified using a Bioo Scientific NEXTprep^TM^-Bacteria DNA Isolation Kit. Libraries were prepared from these strains with the Kapa HyperLibrary Preparation kit (Kapa Biosystems, Wilmington MA), pooled and quantified by qPCR and sequenced for 101 cycles from each of the the fragments on a HiSeq 2500 (Illumina, San Diego, CA). This HiSeq run was performed by the Biotechnology Core Facility at the University of Illinois at Urbana-Champaign and included additional strains not presented here. All other sequencing was performed on a locally operated and maintained Illumina MiSeq system.

For MiSeq runs which generated data for all minimal medium evolved strains and replicates 2 to 4 of the rich medium selection experiments, DNA was extracted with either the Bioo Scientific NEXTprep. kit or the MoBio Ultraclean Microbial DNA isolation kit. Different isolation kits were used due to the discontinuation of the Bioo Scientific kit. DNA was quantified by qubit and Bioanalyzer and libraries were prepared using the NexteraXT kit from Illumina.

Sequencing adapters for the HiSeq generated data were trimmed using *flexbar* (http://sourceforge.net/projects/flexbar/). MiSeq runs were demultiplexed and trimmed using the onboard Illumina software. Analysis was performed using the *breseq* platform (*Deatherage and Barrick, 2014*) in polymorphism mode. *Breseq* uses an empirical error model and a Bayesian variant caller to predict polymorphisms at the nucleotide level. The algorithm uses a threshold on the empirical error estimate (E-value) to call variants (*Barrick and Lenski, 2009*). The value for this threshold used here was 0.01, and at this threshold, with the sequencing coverage for our samples, we report all variants present in the population at a frequency of 0.2 or above (*Barrick and Lenski, 2009*). All other parameters were set to their default values. Reads were aligned to the MG1655 genome (INSDC U00096.3). We note that *breseq* is not well suited to predicting large structural variation. Since we sequence populations at different points during selection, observation of the same mutations at different points in time significantly reduces the probability of false positives (*Lang et al., 2013*).

The founder strain was sequenced at an average depth of 553× when aggregating reads from four separate sequencing reactions. Any mutations observed in this strain were excluded from further analysis. Tables 1,2,3,4,6,7,8,9 document mutations, important mutations were confirmed by Sanger sequencing as noted in the captions to these tables. Since these genomes were sequenced at very high depth, we did not confirm every mutation by Sanger sequencing. All mutation calls made by *breseq* were inspected manually and found to be robust or they were excluded. We also manually inspected the founder strain reads aligned to regions where frequent mutations were observed in the evolved strains (*clpX* E185*, the A1bp mutation at position 523086 and *galS* L22R) to confirm that those mutations were not present in the founder. Sequencing data are available at https://doi.org/10.13012/B2IDB-3958294_V1.

**Table 1.** Rich medium replicate 1: All mutations detected above a frequency of 0.2 in rounds 5, 10 and 15 for rich medium selection replicate 1. The first number in each cell denotes the distance in base pairs from *ori* (location). The second entry (mutation) identifies the mutations with ‘IG’ denoting an intergenic mutation. The third entry (fraction) is the fraction of the population carrying this mutation (as inferred by breseq in polymorphism mode). The fourth entry (coverage) is the number of reads that aligned to this location. In the round 15 strain the *clpX* SNP and ∆1bp deletion at position 523 086 were confirmed by Sanger sequencing.

| Rich medium replicate 1 |  |  |  |
| --- | --- | --- | --- |
| round, (coverage) | 5, (172x) | 10, (213x) | 15, (180x) |
| Mutation (loc., mut., frac., cov.) | 457978, *clpX* E185*, 75.2%, 179<br>523086, Δ 1bp, IG, 100%, 194<br>950518, *pflA* T188I, 22.2%, 144<br>1978458, G→T IG, 21.2%, 156<br>3618863, *nikR* H92H, 20.7%, 189 | 457978, *clpX* E185*, 100%, 199<br>523086, Δ 1bp, IG, 100%, 266<br>990379, A→C IG, 100%, 201 | 457978, *clpX* E185*, 100%, 164<br>523086, Δ 1bp, IG, 100%, 168<br>663115, Δ 1bp *dacA*, 100%, 150<br>990379, A→C IG, 100%, 156 |

**Table 2.** Rich medium replicate 2: All mutations detected in rounds 5,10 and 15 of replicate 2. See Table 1 caption. Note low coverage on Δ 1bp mutation at 523086 noted in **bold**.

| Rich medium replicate 2 |  |  |  |
| --- | --- | --- | --- |
| round, (coverage) | 5, (218x) | 10, (100x) | 15, (166x) |
| Mutation (loc., mut., frac., cov.) | 457978, *clpX* E185*, 100%, 220<br>950518, *pflA* T188I, 27.2%, 210<br>523086, Δ 1bp, IG, 100%, **10/18** | 457978, *clpX* E185*, 100%, 109<br>523086, Δ 1bp, IG, 100%, **16** | 457978, *clpX* E185*, 100%, 184<br>523086, Δ 1bp, IG, 100%, **24**<br>667259, *mrdA* R320H, 39.5%<br>159<br>794472, *modE* L58*, 42.4%, 136 |

**Table 3.** Rich medium replicate 3: All mutations detected in rounds 5, 10 and 15 of replicate 3. See Table 1 caption.

| Rich medium replicate 3 |  |  |  |
| --- | --- | --- | --- |
| round, (coverage) | 5, (291x) | 10, (45x) | 15, (186x) |
| Mutation (loc., mut., frac., cov.) | 457978, *clpX* E185*, 100%, 300<br>523086, Δ 1bp, IG, 50%, **38**<br>950518, *pflA* T188I, 26.3%, 332 | 457978, *clpX* E185*, 100%, 43<br>523086, Δ 1bp, IG, 100%, **8**<br>950518, *pflA* T188I, 53.3%, 53<br>321263 T→C IG, 25%, 16<br>382794 +9b.p. insertion *yaiX*, 64%, NA | 457978, *clpX* E185*, 100%, 185<br>523086, Δ 1bp, IG, 100%, **16**<br>950518, *pflA* T188I, 30.6%, 190<br>1968653, *cheR* Q238K, 29.6%, 190<br>382794 +9b.p. insertion *yaiX*, 25.8%, NA<br>4161562 Δ17b.p. *fabR*, 46.2%, 67 |

**Table 4.** Rich medium replicate 4: All mutations detected in rounds 5, 10 and 15 replicate 4. See Table 1 caption. Note low coverage on Δ 1bp mutation at 523086 noted in **bold**.

| Rich medium replicate 4 |  |  |  |
| --- | --- | --- | --- |
| round, (coverage) | 5, (384x) | 10, (555x) | 15, (333x) |
| Mutation (loc., mut., frac., cov.) | 457978, *clpX* E185*, 100%, 370<br>523086, Δ 1bp, IG, 50%, **72**<br>950518, *pflA* T188I, 31.7%, 446 | 457978, *clpX* E185*, 100%, 559<br>523086, Δ 1bp, IG, 100%, **34/83** | 457978, *clpX* E185*, 100%, 339<br>523086, Δ 1bp, IG, 100%, **19/33**<br>3619915, *rhsBW242G*, 24.9% 20 |

**Table 5.**
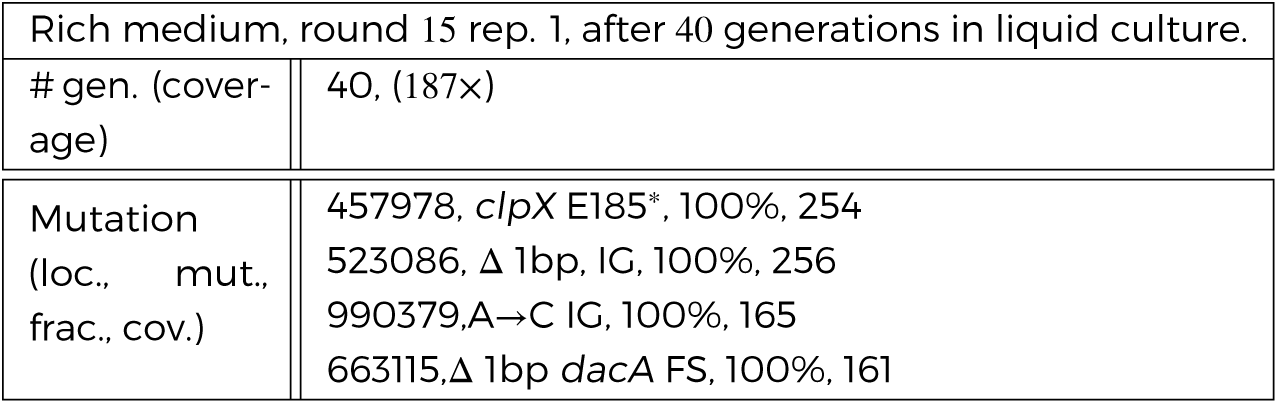
Mutations present after 40 generations of liquid culture growth for rich medium replicate 1 round 15 strain.

**Table 6.**
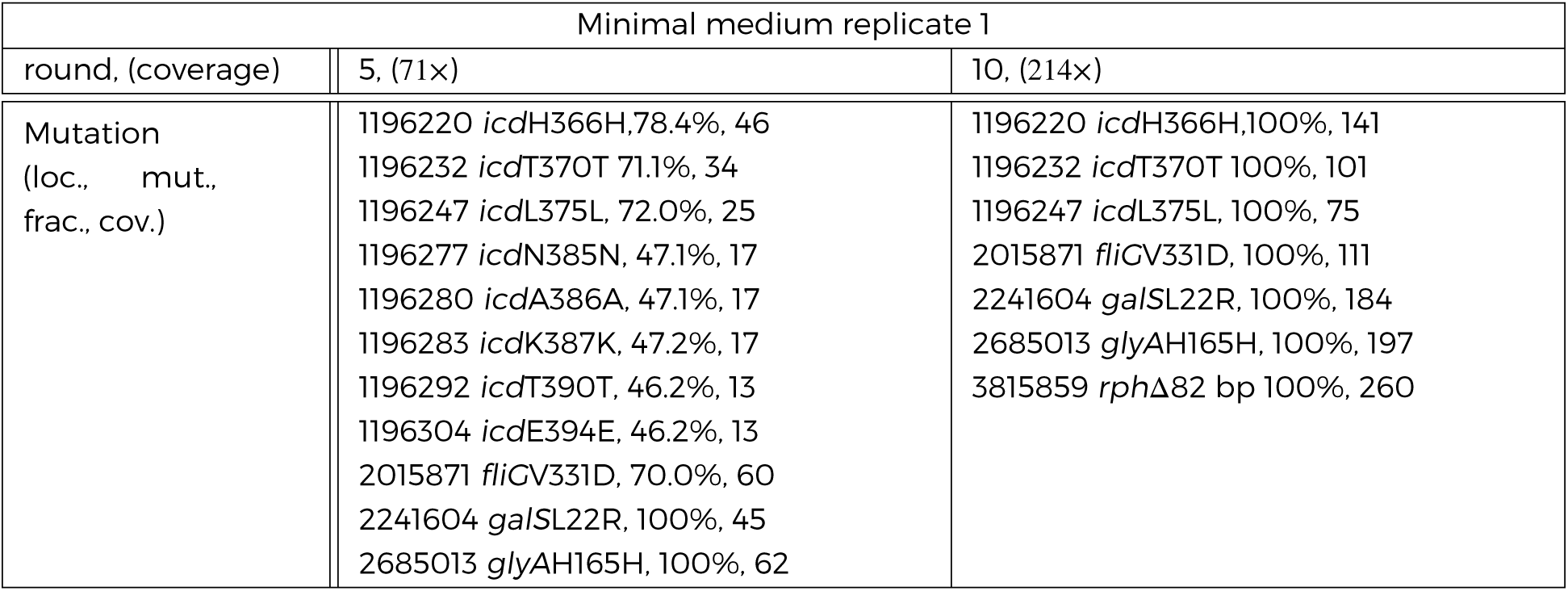
Minimal medium replicate 1: All mutations detected in rounds 5, 10 replicate 1 in minimal medium. The *galS*L22R mutation in rounds 5 amd 10 was confirmed by Sanger sequencing. See Table 1 caption.

**Table 7.**
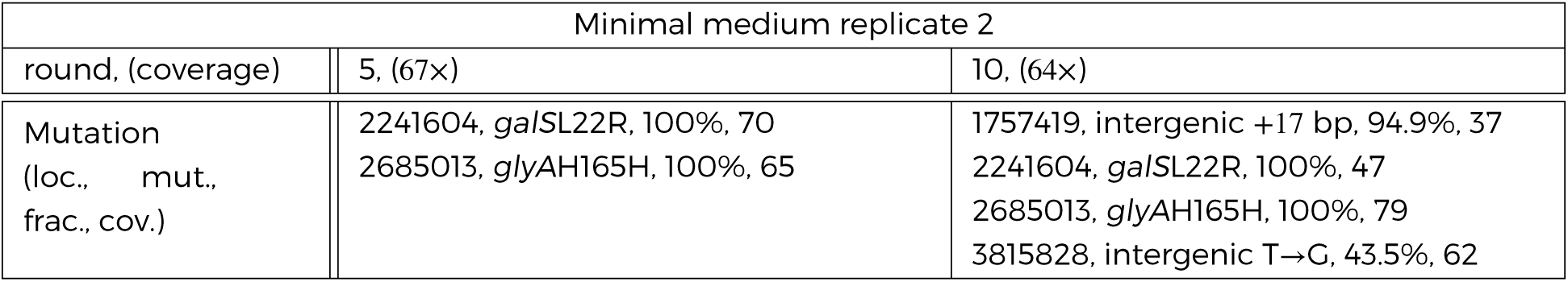
Minimal medium replicate 2: All mutations detected in rounds 5, 10 replicate 2 in minimal medium. The *galS*L22R mutation in rounds 5 amd 10 was confirmed by Sanger sequencing. See Table 1 caption.

**Table 8.**
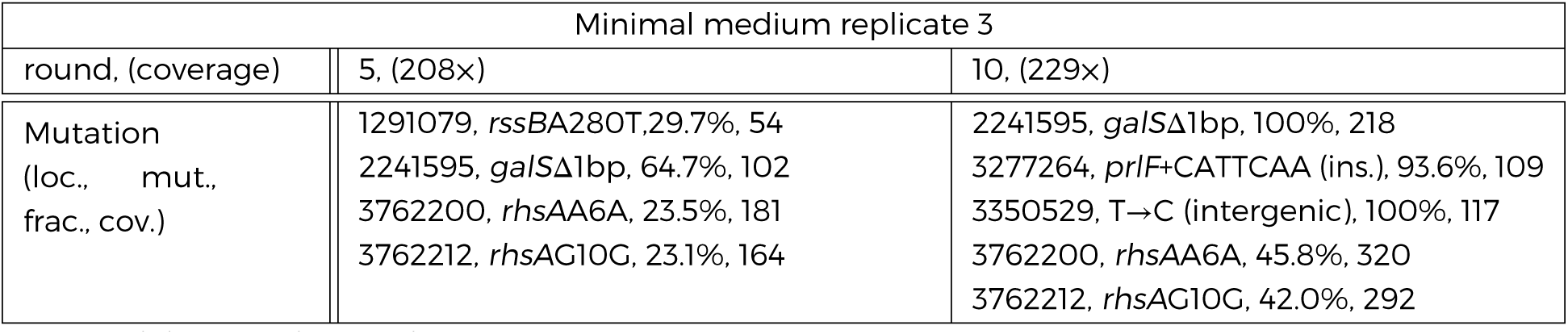
Minimal medium replicate 3: All mutations detected in rounds 5, 10 replicate 3 in minimal medium. See Table 1 caption.

**Table 9.**
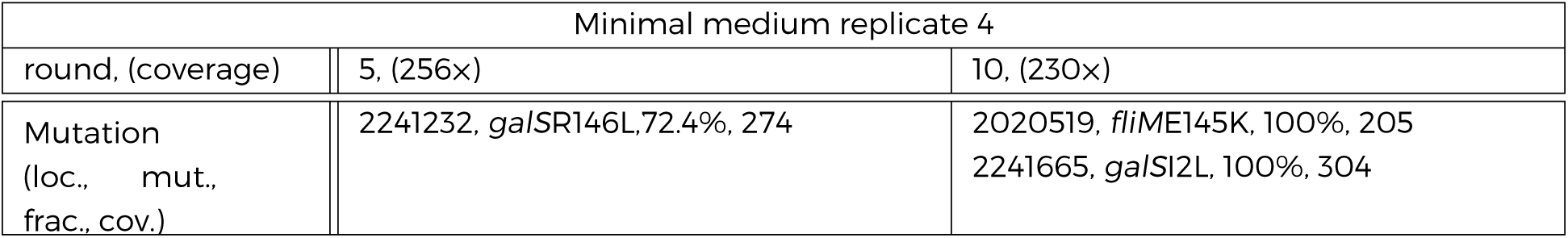
Minimal medium replicate 4: All mutations detected in rounds 5,10 replicate 4 in minimal medium. See Table 1 caption.

**Table 10.** Reaction-diffusion model parameters used in this study

### Mutant reconstruction

Knockout mutants (*ΔclpX, ΔgalS*) were constructed by P1 transduction from KEIO collection mutants (*Baba et al., 2006*). Mutations were confirmed by PCR. Antibiotic markers were not removed prior to phenotyping.

Three commonly single nucleotide polymorphisms (SNPs) observed across evolution experiments were reconstructed in the chromosome of the ancestral background (founder) using a recombineering method presented previously (*Kuhlman and Cox, 2010; Tas et al., 2015*). These mutations were the clpXE185* mutation, the single base pair deletion between *ybbP* and *rhsD* (which we refer to as “Δ1bp”) and galSL22R. For full details of the recombineering we performed see supplementary file 1. Briefly, recombineering proficient cells were prepared by electroporation of the helper plasmid pTKRED (*Kuhlman and Cox, 2010*) and selection on spectinomycin. A linear “landing pad” fragment consisting of *tetA* flanked by I-SceI restriction sites and homologies to the desired target site was synthesized from the template plasmid pTKLP-tetA and site specific primers. The landing pad was inserted by electroporation into recombineering proficient cells and transformants were selected by growth on tetracycline. Successful transformants were confirmed by PCR. A second transformation was then performed using a 70bp oligo containing the desired mutation near the center and flanked by homologies to target the landing pad. Counterselection for successful transformants was performed with NiCl_2_ (6mM for the *ClpX* and *GalS* mutations, 6.5mM for A1bp). Successful recombination at this step resulted in removal of the landing pad and integration of the 70bp oligo containing the desired mutation. The helper plasmid pTKRED was cured by growth at 42°C and confirmed by verifying spectinomycin susceptibility. The presence of desired mutations in the final constructs was confirmed by Sanger sequencing.

## Figure Supplements

Figure 1 - figure supplement 1: Selection with non-chemotactic (ΔcheA-Z) mutant

Figure 1 - figure supplement 2: Change in migration rate during long-term liquid culture

Figure 1 - figure supplement 3: Adaptation in rich medium depends on sampling location Figure 1 - figure supplement 4: Comparison of founding and evolved strains to RP437: Singlecell swimming in rich medium

Figure 1 - figure supplement 5: Persistence of rich medium fast migrating phenotype in rich medium

Figure 2 - figure supplement 1: Reaction-diffusion model recapitulates qualitative features of colony expansion

Figure 2 - figure supplement 2: Comparison of front profiles from simulation and experiment

Figure 2 - figure supplement 3: Simulation of migration rate versus tumble frequency

Figure 3 - figure supplement 1: Microfluidic device and single-cell swimming trajectory

Figure 3 - figure supplement 2: Tumble durations and run lengths for evolved strains

Figure 3 - figure supplement 3: Reproducibility ofthe evolved phenotype

Figure 3 - figure supplement 4: Swimming statistics as a function of culture density

Figure 4 - figure supplement 1: Predicted migration rates for evolved strains

Figure 4 - figure supplement 2: Swimming statistics, growth rates and migration rates for mutants

Figure 6 - figure supplement 1: Determining 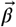 from reaction-diffusion model

Figure 6 - figure supplement 2: Direction of phenotypic evolution with and 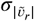 and 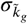

Figure 6 - figure supplement 3: Stochastic simulations of selection in minimal medium

## Source data

Source data file 1

## Supplementary files

Supplementary file 1: additional experimental methods.

## Acknowledgments

We would like to acknowledge Christopher Rao for strains; Yann Chemla, Andrew Ferguson, Hong-yan Shih, Nigel Goldenfeld and faculty members ofthe Center for the Physics of Living Cells for useful discussions. Alvaro Hernandez at the Roy J. Carver Biotechnology Center at the University of Illinois at Urbana-Champaign HiSeq sequencing and Elizabeth Ujhelyi provided assistance with MiSeq sequencing.

## Author contributions

SK conceived of the project. DF, HM, and DS performed the experiments. SK, DF, DS and HM analyzed the data. JM wrote the code to perform numerical simulations ofthe reaction- diffusion model. TEK assisted DF in mutant reconstructions.

